# Reaction Norm Modeling of High-Dimensional Genomic and Environmental Data Improves Prediction Accuracy in Winter Wheat

**DOI:** 10.64898/2026.05.05.722758

**Authors:** Shailesh Raj Acharya, Julian Garcia Abadillo Velasco, Jeanette Lyerly, Gina Brown-Guedira, Diego Jarquin, Nonoy Bandillo

**Affiliations:** Department of Crop and Soil Sciences, North Carolina State University, Raleigh, North Carolina, USA; USDA-ARS SEA, Plant Science Research, Raleigh, North Carolina, USA; Department of Agronomy, University of Florida, Gainesville, Florida, USA

## Abstract

Genomic prediction models that account genotype-by-environment (G×E) have the potential to accelerate the rate of genetic gain for yield and agronomic performance, yet relatively few studies have applied G×E prediction in public soft red winter wheat (*Triticum aestivum*) breeding programs. In this study, we extended a reaction norm-based genomic prediction framework by integrating weather-based environmental covariates to more effectively capture genotype– environment interactions. Key agronomic traits, including seed yield, plant height, test weight, and heading date, were evaluated across 33 environments (location–year) using over 3,200 breeding lines from the North Carolina State University small grains breeding program. Multiple genomic prediction models were compared using several cross-validation (CV) schemes representing common breeding scenarios. Across traits, the reaction norm M5 model, which incorporates both G×E and genotype-by-environmental covariate interactions (G×O), achieved the highest prediction accuracy (PA) in CV2 (predicting incomplete field trials) and CV1 for yield and test weight (predicting new lines). The highest PA was observed for test weight under CV2 (0.54) and for yield under CV1 (0.41). Under CV0 (predicting new environments), the M3 model incorporating G×E produced highest PA across traits, with the greatest accuracy for plant height (0.45), although differences among M2, M3, and M4 were small. Prediction under CV00 (predicting new lines in new environments) remained more challenging, with PA values 0.10 – 0.20 across traits. Overall, our results demonstrate that integrating environmental covariates into genomic prediction models can improve predictive performance across diverse wheat-growing environments in North Carolina, supporting their utility for applied breeding efforts.

**CORE IDEAS:**

- Integrating genotype-by-environment (G×E) interactions with environmental covariates improves prediction accuracy across environments.
- Model performance varies by prediction scenario, with different approaches performing best for new lines, incomplete trials, or new environments.
- Prediction of new lines in new environments remains challenging.

**PLAIN LANGUAGE SUMMARY:** This study explores how adding environmental information to genomic prediction models can improve prediction accuracy in a public winter wheat breeding program. Using data from multi-environment trials conducted across diverse conditions in North Carolina, we evaluated statistical models that capture how different wheat lines respond to changing environments. By incorporating weather data, we improved the ability to predict performance across locations and years. These findings provide practical insights for refining selection strategies and accelerating genetic gain in wheat breeding.

## 1. INTRODUCTION

Soft red winter wheat (Triticum aestivum L.; SRWW) constitutes the largest class of wheat cultivated in the southeastern United States. Of the 342 million bushels produced in this region, approximately 18 million bushels are grown in North Carolina (NC), contributing an estimated $100 million to the state’s economy and serving as a key driver of NC’s baking and milling industries (USDA, 2025). Wheat is grown across diverse regions of NC that span multiple environments. This environmental heterogeneity presents a major challenge for breeders aiming to develop high-yielding, broadly adapted cultivars. Consequently, understanding genotype-by-environment (G×E) interactions is essential for identifying wheat lines that combine high yield potential with stable performance across environments.

In wheat breeding programs, strong G×E interactions imply that genetic merit cannot be accurately assessed without accounting for environmental variation among testing sites. Prediction models that ignore G×E and relevant environmental covariates often fail to capture true genotype performance (Crossa et al., 2017). Reaction norm models address this limitation by extending genomic best linear unbiased prediction (GBLUP) to incorporate environmental covariates into genomic effects, enabling more reliable predictions across years and locations (Jarquín et al., 2014). Within this framework, genotypic performance is modeled as a function of environmental gradients, allowing prediction in both observed and unobserved environments while capturing genotype-specific sensitivities to environmental variation (Monteverde et al., 2019).

Incorporating weather-derived environmental covariates such as temperature, precipitation, solar radiation, and growing degree days further enhances predictive accuracy (Morais Júnior et al., 2018; Raffo et al., 2022). When summarized over key developmental stages, these variables better reflect the conditions experienced by crops and improve alignment between environmental inputs and physiological responses. Embedding such covariates into covariance structures or kernel-based models enables more precise partitioning of environmental similarity and genotype-specific responses (Jarquín et al., 2021). As a result, this approach not only improves prediction across diverse and future climates but also provides insights into the environmental drivers of trait expression, supporting more informed selection and climate-resilient breeding strategies.

However, the practical value of incorporating weather covariates ultimately depends on their ability to improve prediction accuracy under realistic breeding scenarios. In multi-environment trials, the primary objective is not only to explain past performance but also to predict the performance of untested genotypes in new or sparsely sampled environments. The benefit of environmental covariates may therefore depend on the amount of available phenotypic data and on how effectively these variables capture meaningful environmental differences. Previous studies have shown that while modeling G×E generally improves prediction accuracy, the inclusion of weather covariates does not always lead to increased accuracy and is highly dependent on how environmental information is represented (Jarquín et al., 2017, 2021).

To rigorously assess their utility, genomic prediction models must be evaluated under cross-validation (CV) schemes that reflect real-world breeding decisions. Commonly used strategies include CV1, which evaluates prediction of new lines in previously tested environments, and CV2, which assesses performance for lines evaluated in only a subset of environments (Jarquín et al., 2020; Monteverde et al., 2019; Sukumaran et al., 2018). More recent approaches extend these frameworks to include prediction in entirely new environments (CV0) and forward prediction into future years, leveraging historical data to simulate realistic forecasting scenarios (McBreen et al., 2025). The most stringent scenario, CV00, evaluates the ability to predict new genotypes in completely unobserved environments.

In this study, we evaluated genomic prediction accuracy for four key traits including grain yield (Y), test weight (TW), heading date (HD), and plant height (HT) in SRWW from the North Carolina State University small grains breeding program. Using a reaction norm framework, we compared five prediction models across four cross-validation schemes (CV1, CV2, CV0, and CV00) that represent distinct breeding scenarios. The dataset comprised lines evaluated across 33 environments (location–year combinations) from 2008 to 2023, incorporating five weather variables obtained from NASA. Our objective was to determine whether incorporating G×E interactions and weather covariates improves prediction accuracy under practical breeding conditions, including the prediction of untested genotypes, incomplete field evaluations, novel environments, and future growing seasons.

## 2. MATERIALS AND METHODS

### 2.1 Plant phenotypic data

Data for four traits: grain yield (Y), test weight (TW), heading date (HD) and plant height (HT) were obtained from NCSU soft red winter wheat lines grown across eastern, central and western North Carolina from 2008-2023 consisting of a total of 3196 breeding lines. The breeding lines consisted of a diverse range of lines from preliminary (NCOBS), advanced (NCWAT), and legacy (NCLEGACY) sources. Additionally, lines from regional nurseries that originated from various breeding programs of the SunGrains breeding collaborative in the southeastern United States, including GAWN and Sunwheat, as well as released check varieties were included.

Yield was observed across all 33 environments with the minimum genotypes overlap in CLAYTON2018 and maximum genotypes overlap in KINSTON2021. The total number of environments and their respective overlap among genotypes across these environments for grain yield is shown in Figure 1.

**Figure 1:**
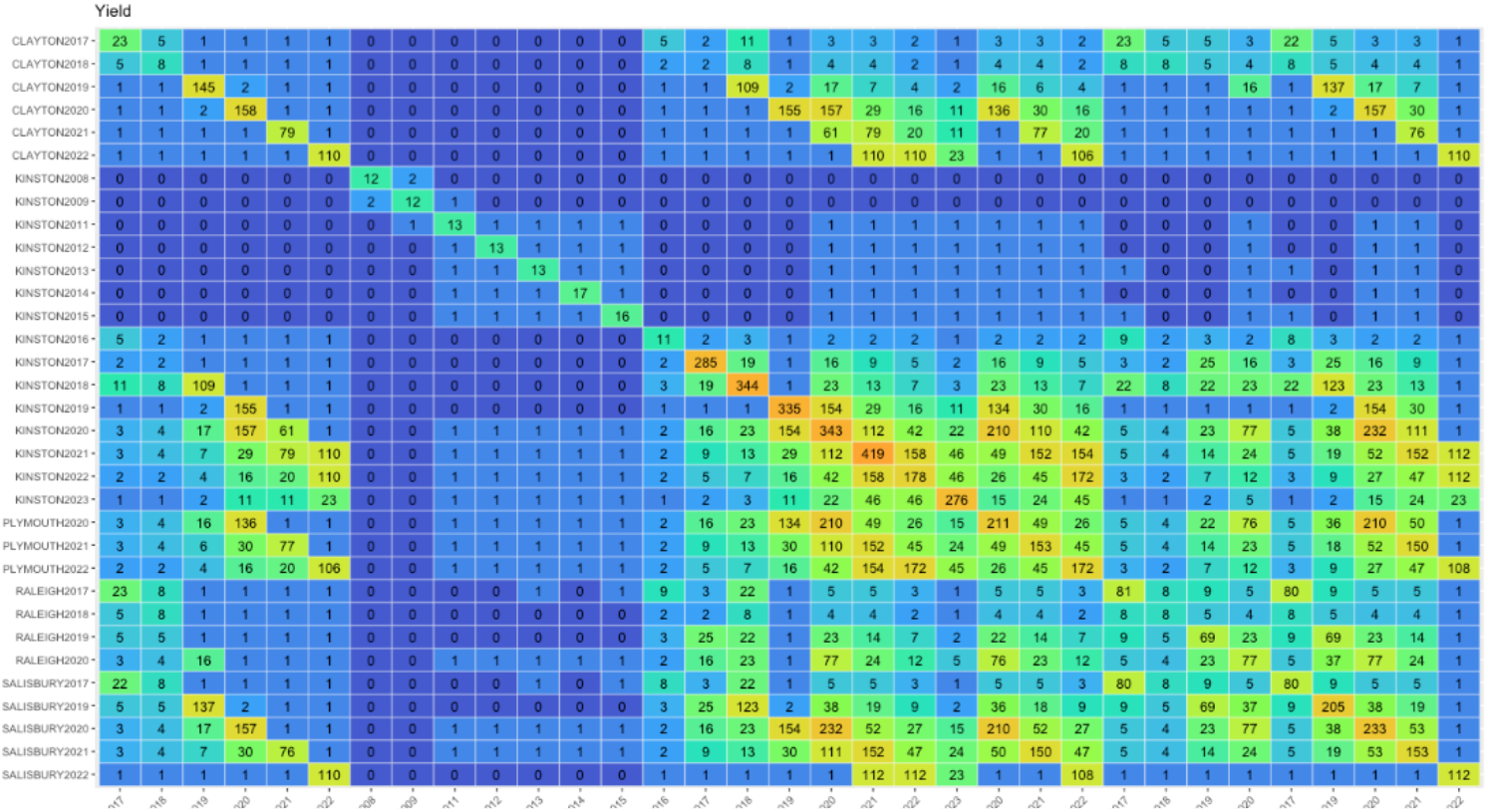
Total number of overlapping genotypes between environments for ‘Yield’. Yield was observed across all 33 environments. The 33×33 matrix shows the number of lines in each environment (location-year combinations) on the diagonal. The upper off-diagonal shows common lines between environments, while the lower off-diagonal shows lines not shared between environments.

Similarly, the total number of environments tested for all remaining traits: test weight, heading date and plant height is presented in Table 1, and their environment specific overlap in a similar matrix form is present in Figure S1.

**Table1:** Total number of environments tested and their mean and median values across all four traits for 3196 wheat breeding lines from 2008-2023.

| Trait | Environment tested | Min.Overlap | Max.Overlap | Mean Genotype Number | Median Genotype Number |
| --- | --- | --- | --- | --- | --- |
| Yield (bu/ac) | 33 | 8 | 419 | 132.2 | 110 |
| Test Weight (lb/bu) | 33 | 8 | 478 | 143.3 | 111 |
| Heading Date (Jul) | 14 | 8 | 648 | 214.7 | 84 |
| Plant Height(in) | 11 | 8 | 145 | 48.8 | 23 |

### 2.2 Genotypic data

DNA extraction, genotype-by-sequencing (GBS) library preparation and SNP calling were conducted as previously described by (Winn et al., 2023). High quality SNP data was obtained after removing genotypes with 85% or more missing data, filtering out markers with over 50% missing data, minor allele frequencies less than 0.05 and 10% heterozygosity using TASSEL v5. Thereafter, SNP imputation to account for missing data was performed using Beagle version 5.2 (Browning & Browning, 2016). Ultimately, a total of 40,850 high quality SNP markers were retained for downstream modelling.

### 2.3 Weather data

Daily weather variables including minimum and maximum temperature, precipitation and relative humidity were obtained from the NASA POWER (Sparks, 2018) database. Daylength (photoperiod) in hours was calculated from the latitude and day of year using a standard trigonometric approach implemented in R with lubridate (Grolemund & Wickham, 2011). The time interval for a year started from 1^st^ of October to 1^st^ of September of the following year (For example data for 2022 consists of dates between Oct-1-2022 to Aug-31-2023). Data was then averaged by month so that the total number of features per environment was the product of the five weather variables and 11 months, resulting in 55 weather features per environment. The geographic coordinates of all sites from which weather data was extracted are presented in Table S2.

### 2.4 Statistical analyses of phenotypic traits

Genotypic best linear unbiased estimates (BLUEs), which represent adjusted phenotypic values for downstream modelling, were obtained for all traits: grain yield (Y), test weight (TW), heading date (HD) and plant height (HT) using ASReml-R v4.2 (Butler et al., 2023).

Each phenotypic trait for genotype i, in replication j, within environment e was modeled as:

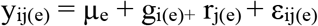

where µ_e_ is the fixed mean effect of environment (e), y_ij_ is the observed phenotypic value of i^th^ genotype in j^th^ replication of e^th^ environment, g_i(e)_ is fixed genotype effect of i^th^ genotype in e^th^ environment, r_j(e)_ is the random replication effect within environment (e), and ε_ij(e)_ is the residual which is assumed to be independently and identically distributed (IID).

Broad sense heritability (H^2^) was estimated separately for each environment across all traits using the linear mixed model described above, but with genotype and replication effect specified as random.

Broad sense heritability (H^2^) was calculated as:

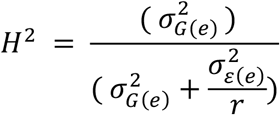

where 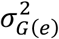 and 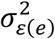 correspond to variance due to genotypic and residual effects and r represents the number of replications.

### 2.5 Prediction Models

Five different models were used to predict the performance of all four phenotypic traits based on foundational model structure described by (Jarquín et al., 2014). BLUEs obtained from phenotypic traits served as the input of all these models and all factors in the model were treated as random effects. A schematic diagram of all five models is presented in Figure 2.

**Figure 2:**
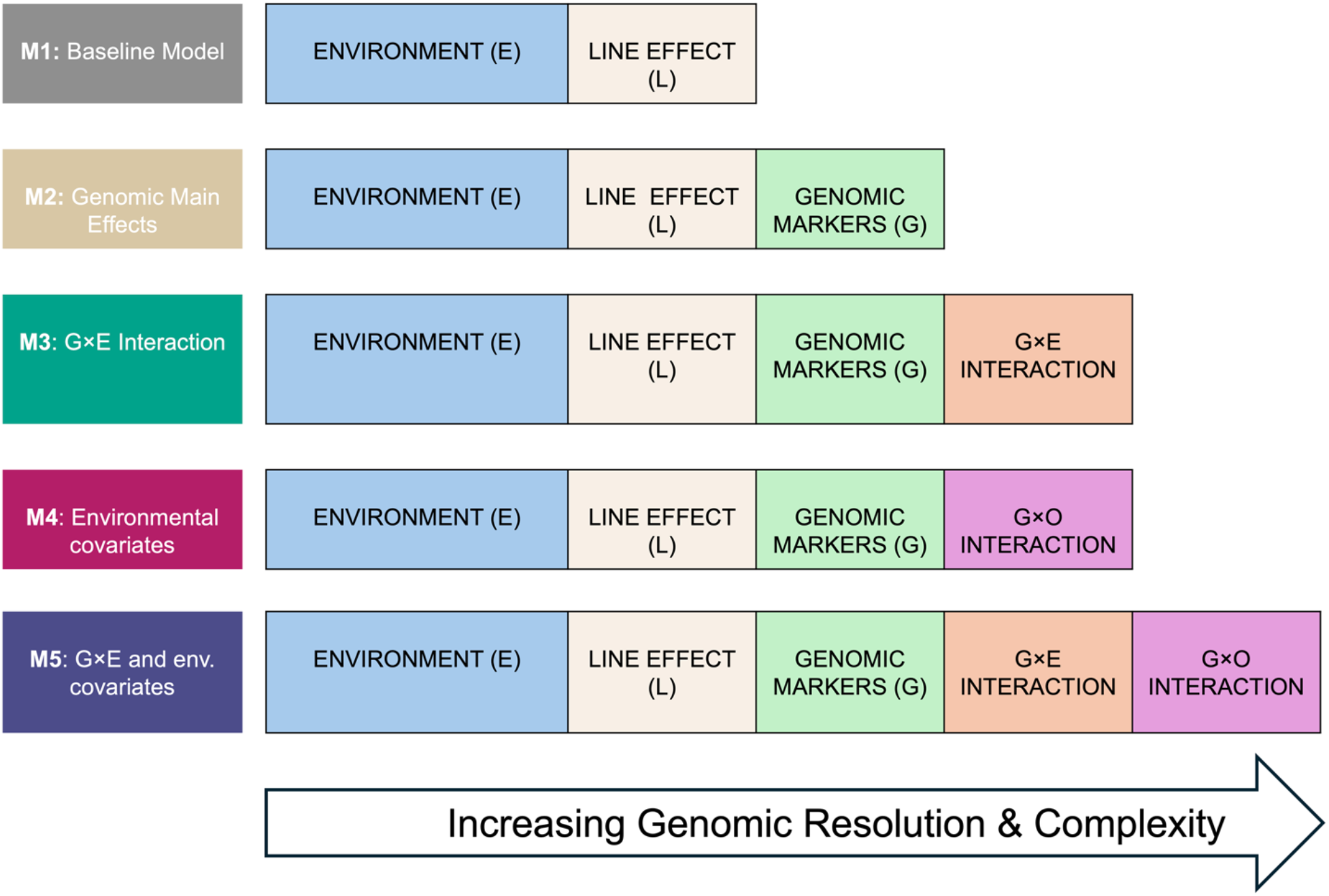
Schematics of all five different prediction models. Model complexity increases as more variables are added, which can also improve prediction accuracy and enhance genomic resolution.

#### 2.5.1 M1: Baseline model (E + L)

Model M1, also called baseline model, integrates main effects for both environment (E) and line (L) to predict traits and is described as:

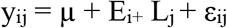

The model contains overall mean (µ), random i^th^ environmental effects (E_i_) and j^th^ line effect (L_j_) assuming:

E_i_ ~ N (0, σ^2^_E_) with IID, where σ^2^_E_ is the variance component of environment,

L_j_ ~ N (0, σ^2^_L_) with IID, where σ^2^_L_ is the variance component of line, and

ε_ij_~ N (0, σ^2^ε) with IID, where σ^2^ε is the residual variance.

As both E and L assume independence across different levels of environment and line, this strong dependency assumption makes the model unable to borrow information from related environments/genotypes. In practice, if a line or environment has not been observed, then this model becomes null as it just predicts the average (µ). Hence, it was used as a negative control to quantify the differences between CV2 (random cross-validation) and CV1(genotype constrained cross validation).

#### 2.5.2 M2: E +L+ G

In the M2 model, the SNP markers were introduced in the baseline model as:

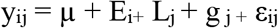

where g _j_ follows multivariate normal distribution where g = {g_o_} ~ N (0, Gσ^2^_g_). The covariance matrix Cov(g) = Gσ^2^_g_, where 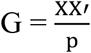 denotes genomic relationship matrix, X is the centered and standardized genotype matrix to *p* SNP markers, and σ^2^_g_ is the corresponding genomic variance.

The G term in the M2 model accounts for genotypic effects where SNP marker information is used to establish relationships between each pair of genotypes. This allows borrowing information such that if a given genotype performs well, then a related genotype is expected to perform in a similar manner.

#### 2.5.3 M3: E +L + G + GE

Model M3 extends model M2 and includes a × interaction component.

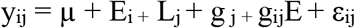

The model is similar to M2 with the addition of g_ij_E component, which denotes interaction of i^th^ genotype with j^th^ environment with gE = {g_ij_E}~ MVN (0, (Z_g_GZ^T^_g_) ° (Z_E_ Z^T^_E_) σ^2^_Eg_) where Z_g_ and Z_E_ are the incidence matrices for lines and environment respectively. σ^2^_Eg_ is the corresponding variance component of g_ij_E. The sign ‘°’ is the Hadamard product between two matrices that denotes elementwise product between two matrices.

The GE term accounts for interaction between genotype and environment allowing different behaviors of the same genotype in different environments. This enables model adaptation and superiority.

#### 2.5.4 M4: E +L + G + GO

In model M3, it was assumed that environments were independent for GE interaction purposes, i.e although two related genotypes will perform similarly in a given environment, it is impossible to predict whether two environments will show the same patterns. The GO term uses weather data to define ‘mega-environments’, where genotype-by-environment patterns are assumed to be similar. Model M4 is:

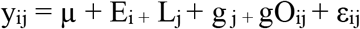

where, gO = {gO_ij_} ~ MVN (0, (ZgGZTg) ° (O O^T^) σ^2^_gO_) where gO_ij_ is the effect of i^th^ genotype at j^th^ environment based on its reaction to weather covariate.

In this equation O O^T^ replaces Z_E_ Z^T^_E_ as a measure of similarity between environments based on covariates. σ^2^_GO_ is the variance of GO interaction.

As we were interested in within-environment variation, the O term is not included in this model because it is a main effect that applies to every phenotype in a given environment, and therefore captures only the average effect of weather across all genotypes.

#### 2.5.5 M5: E +L + G + GE + GO

Model M5 integrates all different terms introduced from M1 to M4 and is the most complex model among all models.

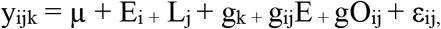

All previously described terms from model M3 and M4 are contained in M5 with both GE and GO interaction present in this model.

This model combines both GE and GO to approximate the ranking within a given environment under the assumption of environmental relatedness according to weather data (GO term) and assumption of unrelatedness (GE term). While GE captures how genotypes react differently to discrete environments, GO captures plasticity of genotypes across environmental gradients.

### 2.6 Variance component estimation

Variance components were estimated to partition genetic background, marker based genetic effects, and environment and weather dependent genetic responses while accounting for residual variability.

The total phenotypic variance was defined as:

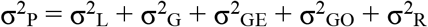

where, σ^2^_L_ is the line variance, σ^2^_G_ is marker based genomic variance, σ^2^_GE_ is genotype × environment interaction variance, σ^2^_GO_ is genotype × weather covariate interaction variance and σ^2^_R_ is residual variance.

### 2.7 Cross validation

To evaluate accuracy of all five implemented models, cross-validation (CV) schemes were implemented based on the strategy described by (Jarquín et al., 2017). To validate models, the phenotypic dataset for all four traits were split into different training and testing sets using k-fold cross validation schemes.

#### 2.7.1 CV2: Prediction of lines in incomplete trials

This cross-validation scheme is used to predict known genotype performance in environments where some data are missing for certain genotypes. Lines were randomly assigned to five cross-validation folds, and the process was repeated 10 times. In each fold, 80% of phenotypic trait data were used for training and 20% for validation. For training and testing split, it was ensured that the training set may include observation of same the genotype in other environments (Eg: Line 1 in Env.2 or Line 2 in Env.1), but never the target genotype in the target environment (E.g. Line1 in Env.1) since that would constitute data leak. Following cross-validation, prediction accuracy was summarized using mean and standard error.

#### 2.7.2 CV1: Prediction of new lines in known environments

CV1 scheme predicts new lines that have not been observed in previously tested environments. Since the fold in CV1 is constrained by genotypes, whole genotypes were held out during cross validation rather than individual observations. Thus, CV1 scheme evaluates if the model can predict new genotypes based on genetically related genotypes it has already seen.

Data for a given genotype were held together in same fold such that none of them are used for training when that genotype is being predicted. Data from CV1 was also assigned to five-folds with 80-20 % training-testing split and the process was repeated 10 times. Following cross-validation, correlation between observed and predicted values within the same environments were calculated.

#### 2.7.3 CV0: Prediction of lines in untested environments

CV0 scheme aims to predict performance of lines in previously untested environments. While CV1 was constrained by genotypes, CV0 is constrained by a new environment. Thus, CV0 aims to predict genotype performance in those environments that were not used in model training but were previously observed in other environments. In the CV0 scheme, the testing environment was completely left out, and the model was trained using data from all remaining environments. This trained model was then used to predict line performance in the earlier excluded environments. Since environments were excluded one at a time, CV0 does not involve random partitioning like CV1 or CV2 and thus was performed only once.

#### 2.7.4 CV00: Prediction of untested lines in untested environments

CV00 scheme combines both CV1 and CV0 restrictions and aims to predict new lines in new environments. In CV00 strategy, the model was trained using untested lines that are predicted in environments where they have not been evaluated using information from different sets of lines tested in other environments. The model was trained using data from remaining genotypes evaluated in other environments. Since CV00 scheme involves systematic exclusion of both genotypes and environments rather than random sampling, CV00 was performed only once. Figure 3 summarizes the cross-validation schemes implemented in this study.

**Figure 3:**
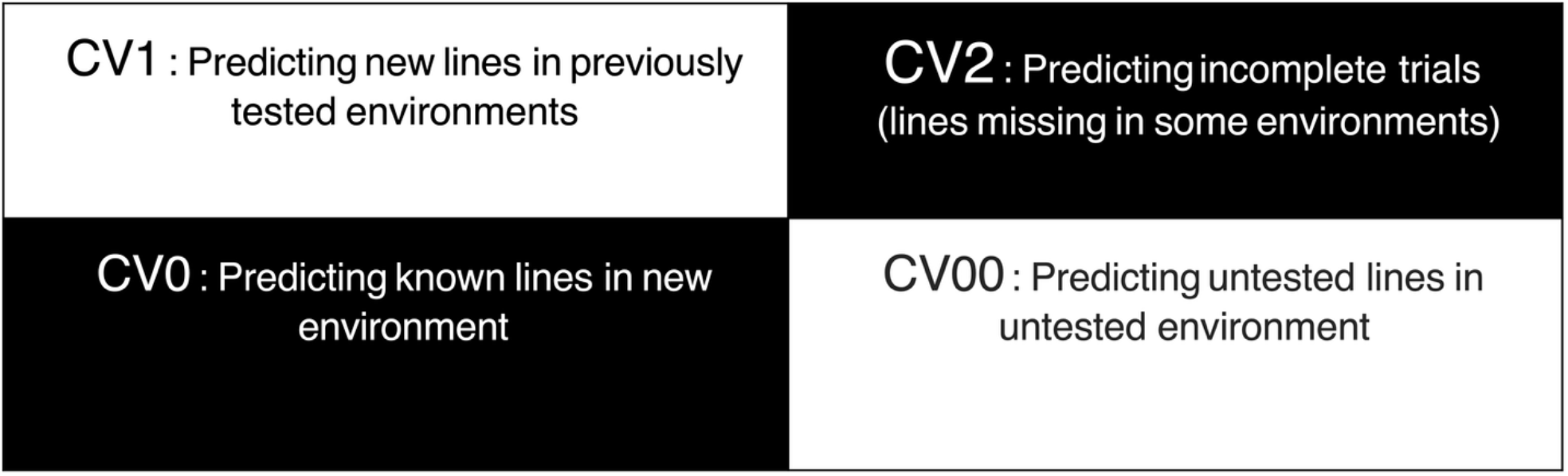
Schematics of four cross validation schemes to evaluate prediction accuracy of five different models (M1 – M5).

### 2.8 Prediction accuracy

The prediction accuracy of all five models tested across four cross validation schemes was obtained by calculating correlation between observed and predicted values. The values from different environments were averaged using the number of observations that led to the correlation as the weight factor. The overall accuracy was then reported as:

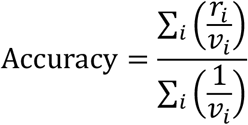

v_i_ represents the weight factor where:

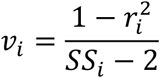

r_i_ represents mean prediction accuracy for each environment and SS_i_ represents total number of genotypes for that environment.

## 3. RESULTS

### 3.1 Phenotypic results

The phenotypic dataset showed that grain yield and test weight had the highest number of testing locations (33) while plant height had the fewest followed by heading date (Table 1). This is typical in a public breeding program where yield-related traits are prioritized more over agronomic traits because of limited resources for gathering data on multiple traits. However, across all 33 environments, heading date had the maximum number of genotypes overlap (648) followed by yield and test weight, likely because agronomic traits like heading time are easier and quicker to measure when assessed across multiple environments. Table 1 summarizes mean and median genotype numbers for all four traits measured across testing environments.

Due to the low level of genotypes overlapping, it is challenging to compute genetic correlations between environments for a given trait. However, we can pair observations of the same genotypes for different traits. In doing so, significant correlation was found for yield – heading date and test weight – plant height pairs. (Figure S2).

Heritability (H^2^) varied across environments and traits with heading date showing highest H^2^ value ranging from 0.35 - 0.95, while test weight had the highest in the CLAYTON 2022 environment reflecting environmental influence on expression of phenotype. H^2^ for yield ranged between 0.33 - 0.87 (Table S1)

### 3.2 Weather data correlation

Five weather variables obtained from NASA POWER were used to construct a kinship matrix, which was then clustered using agglomerative hierarchical clustering (Figure 4). Strong associations of weather variable were observed within the same year compared with across years regardless of locations. This pattern is expected as weather conditions within a given year tend to be more similar than those of past or future years making weather an important factor to consider in breeding cycle decisions. Furthermore, principal component analysis (PCA) was done where environments were clustered by year. Two environments from the same year were more similar than from two environments from the same location across different years. The first two principal components (PC) explained 46.03% and 26.71% of the variance respectively. A total of 80% and 95% of the variance was explained with 3 and 5 PCs respectively (Figure S3).

**Figure 4:**
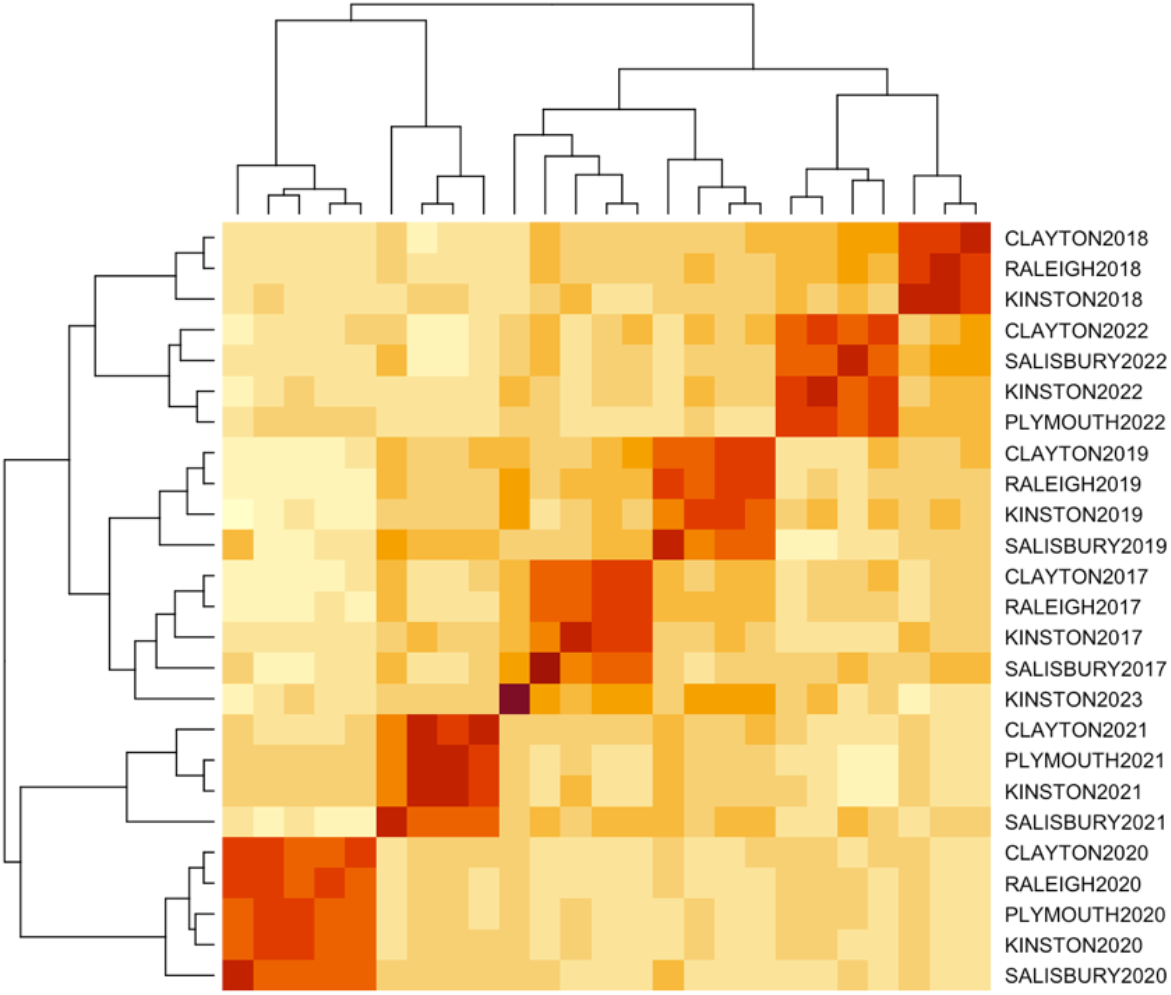
Environment variance-covariance matrix derived from weather covariables and clustered using the agglomerative hierarchical clustering algorithm. Distinct clusters highlighted in red squares show that weather data from different environments within the same year were more closely associated than data from different years, regardless of location.

### 3.3 Variance component analysis

Phenotypic variance (before cross-validation) explained the relative contribution of each term (E, G and O) and their interactions (G×E and G×O) across all four traits. The percentage of variance explained by line (varL) in the baseline model (M1) can be seen as broad-sense heritability while the variance due to marker (var G) is more aligned with narrow-sense heritability. Residual variance (var R) represented variance that could not be explained by any of the terms in the model.

Variance components were estimated both with and without inclusion of environment as main effects to fully describe variance partitioning. As prediction accuracy in our study was evaluated based on within environments, and within-environment variance reflects variance relevant for comparing genotypes within trials, variance partitioning was calculated after accounting for environmental means (i.e excluding varE) and is presented in Figure 5. The complete variance partitioning including environmental variance (varE) is included in Table S3.

**Figure 5:**
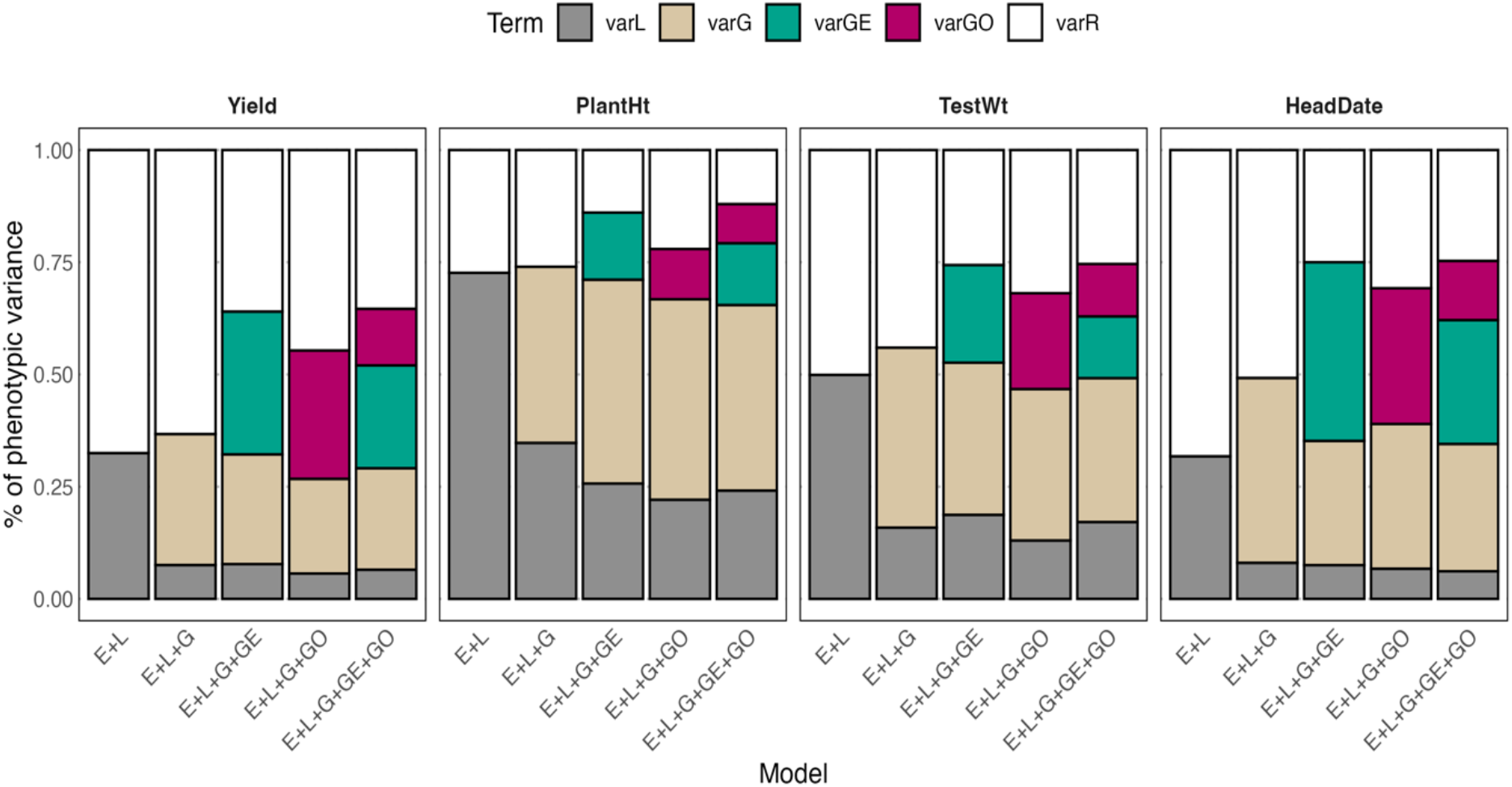
Partitioning of phenotypic variance measured across all four traits. The percentage of variance explained by each term other than environment is shown in the y-axis. Models are denoted in the x-axis, and each trait is represented in a different set of columns. varL represents variance due to line/genotype, varG represents variance due to SNP marker, varR represents residual variance, varGE and var GO represents variance due to interaction terms G ×E and G ×O respectively.

In the baseline model M1 (E+L), phenotypic variance was partitioned into varL and varR. For three traits: yield, test weight and heading date, the largest proportion of phenotypic variance was due to varR indicating substantial unexplained variability beyond genetic differences among lines (Figure 5). However, for plant height, varL exceeded varR indicating strong genetic control across environments. With the addition of G in the second model (M2), the sum of L + G could explain slightly more variability than L alone in the baseline model. Subsequently, as additional term (G×E) was added to M3 model, the genotype-by-environment interaction variance (var GE) explained more variance by further reducing residual variance. In the fifth model (M5) where all terms were combined, residual variances were like those of M3 model, suggesting that GE alone captured genotype-by-environment interaction variance and GO term might be somehow redundant.

### 3.4 Model evaluation using cross-validation

Prediction accuracy (PA) was evaluated within environments as the correlation between observed and predicted values. The overall accuracy obtained as the weighted average of PA for all four traits are represented in Figure 6 and Table S4.

**Figure 6:**
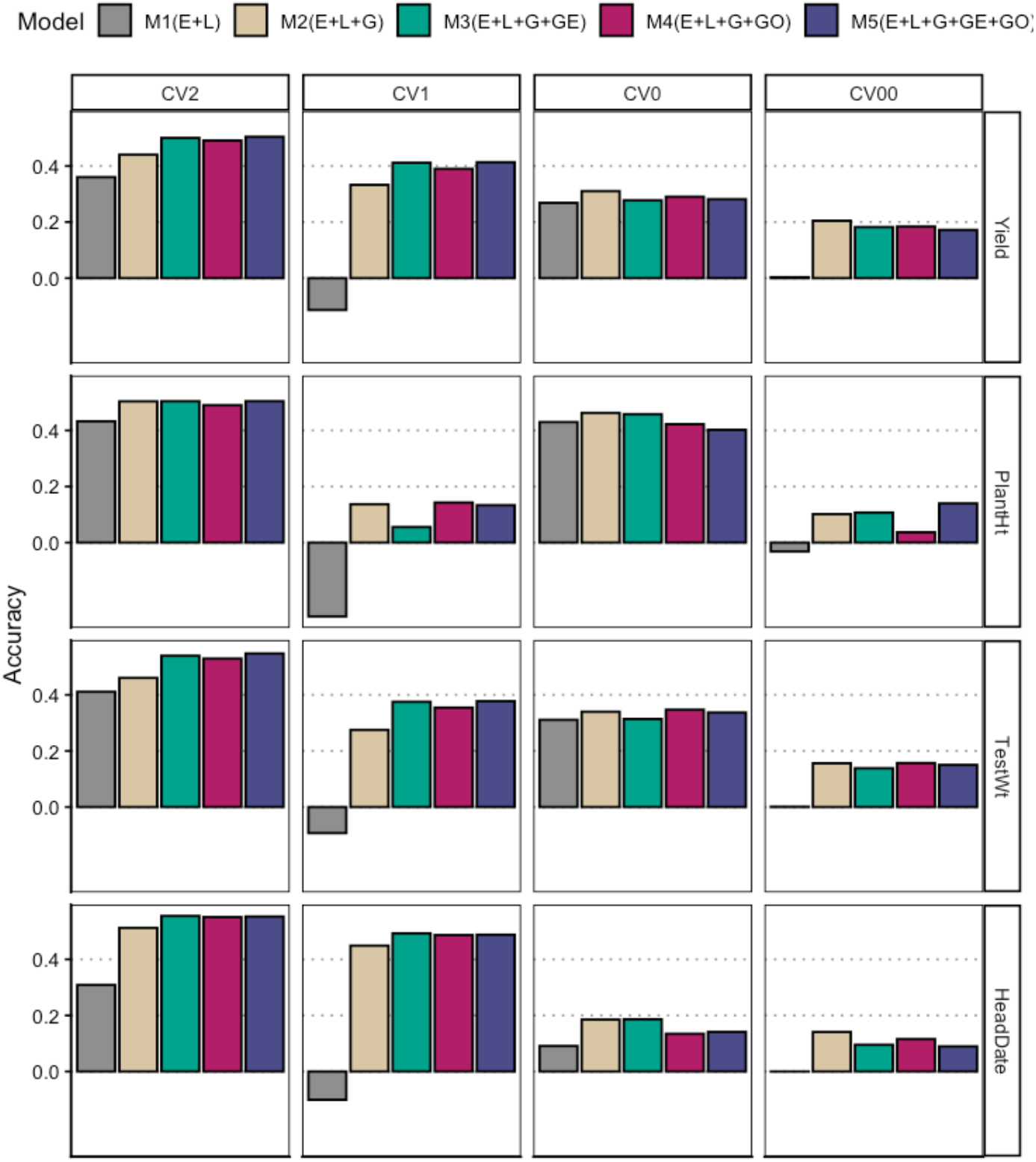
Accuracy achieved by all five models (M1-M5) under various cross-validation scenarios (columns) across four traits (rows). In each grid, accuracy is indicated on the y-axis, while models are represented on the x-axis, corresponding to their respective colors.

#### 3.4.1 Predicting lines in incomplete trials (CV2)

The average accuracy values in predicting lines in incomplete trial was highest for model M5 which includes full reaction norm model including GE and GO. Highest accuracy values across yield (0.50), plant height (0.50) and test weight (0.54) suggests that inclusion of genotype-by-weather covariate provided accuracy gain over genotype-by-environment alone. Meanwhile, model M3 had highest accuracy for heading date (0.55). Although overall highest accuracies were observed for M5 models, the differences between M3 and M5 model were small.

#### 3.4.2 Predicting new lines (CV1)

The model M5 exhibited the highest accuracy in predicting new lines within a known environment, particularly in yield (0.41) and test weight (0.37). However, the differences between M5 and M3 predictions were marginal for both yield and test weight. Notably, these accuracies were 18% lower in yield and 31% lower in test weight compared to the predictions observed in the CV2 scenario.

In contrast, the M3 model demonstrated the highest accuracy in predicting plant height (0.14) and heading date (0.493). However, these accuracies were lower in comparison to the predictions observed in the CV2 scheme.

#### 3.4.3 Predicting new environment (CV0)

Based on the prediction of new environment from known genotype information using CV0 scheme, all five models (M1-M5) produced similar prediction accuracies. While accuracies differed among four traits, with plant height showing highest overall accuracy (0.46), and heading date showing lowest (0.18), differences in accuracies among models within each trait were minimal.

#### 3.4.4 Predicting new lines in new environment (CV00)

In the CV00 scenario, prediction accuracy was consistently 0.2 and below, across all four traits indicating poor predictive performance for new genotypes in unobserved environments. Given the increased complexity of CV00 prediction scenario, under a new environment, it might be better to predict an average performance across environments rather than trying to interpolate adaptation patterns using weather data.

Following prediction accuracy, mean squared error (MSE) was computed as the average of the squared differences between observed and predicted values. The MSE values were similar within the pairs CV2-CV1 and CV0-CV00, with lower MSE observed for CV2-CV1 than for CV0- CV00. The relatively low MSE values for CV2 and CV1 in yield, test weight and heading date indicate that the addition of interaction terms (GE and GO) in models M3-M5 resulted in better predictive performance (Figure S4). Meanwhile, for plant height, model M4 showed the lowest MSE in CV1 relative to CV2. In CV0, model M5 achieved the lowest MSE for yield, model M3 for plant height and M4 model for test weight. For heading date, the baseline M1 model had the lowest MSE, suggesting that the inclusion of SNP markers and weather variates did not improve predictive performance. In CV00, model M2 achieved the lowest MSE for yield, test weight and heading date, whereas M3 and M5 models produced significantly lower MSE values for plant height, suggesting that plant height is strongly influenced by G×E interactions.

## 4. DISCUSSION

The NC wheat growing region encompasses a diverse range of environments spanning across the state, resulting in substantial environmental variability. Thus, modeling multiple traits across multiple environments is essential to improve prediction accuracy when selecting high performing or superior lines in wheat breeding programs. Accounting for year-to-year variation across such multi-environment scenarios requires modeling genotype-by-environment (G×E) interactions to improve trait prediction, particularly grain yield, while other agronomic traits such as plant height and heading date, remain equally important for identifying superior genotypes.

Because these phenotypic traits vary across planting locations and years as different wheat genotypes experience different environmental conditions, genomic prediction in multi-environment trials (MET) must account for G×E interactions to achieve accurate predictions. In our study, we partitioned the predictable variation into main genetic and G×E interactions and then incorporated environmental covariates and their interactions with genotypes (G×O) to assess their contribution to predictive performance. Across cross-validation (CV) scenarios that mimic real-life breeding situations, models that included environmental effects and covariate interactions with genotypes improved prediction accuracy. These findings are consistent with the reaction norm model (RNM) framework which captures G×E by allowing genotypes performances to vary across environments while borrowing information across similar environments. Recent studies have similarly used the RNM framework in METs and demonstrated improved prediction accuracy particularly when incorporating environmental covariates (Brault et al., 2025; Han et al., 2025; Jarquín et al., 2017; Li et al., 2021) and, in some cases, high throughput phenotyping data (McBreen et al., 2025). Accordingly, the application of five RNM-based models (M1-M5) in our study enabled the evaluation of predictive performance for four traits across different CV schemes.

In the first scenario, the CV2 scheme which represented incomplete trials, showed that the full reaction framework (M5), including both G×E and G×O interaction, improved prediction accuracy for yield, plant height and test weight. This suggests that these traits show consistent responses across environments and that genotypes may differ in their sensitivity to weather beyond G×E. Similarly, Jarquín et al. (2014) reported that incorporating environment covariates into the RNM substantially improved wheat yield prediction and explained a meaningful proportion of the variance among wheat lines.

In the second scenario CV1, which predicts new untested lines, the full M5 model gave the best predictions for yield and test weight because the weather covariates probably helped characterize difference among environments. The G×O interaction further allowed lines to differ in their responses to environmental conditions described by these covariates even when the predicted new lines had no phenotypic data. By contrast, under both CV1 and CV2, heading date was best predicted by model M3, which included only G×E, indicating that environment covariates did not add any useful information. For a trait such as heading date, which is often controlled by stable timing cues such as temperature and daylength already captured by location-year effect, G×E may be sufficient. Consistent with our findings, a similar study (Puglisi et al., 2026) also reported high prediction accuracy for heading date under comparable CV schemes based on reaction norm model framework

In the third scenario CV0, model M2 outperformed the other models for all traits. Because this scenario involves predicting lines in unobserved environment without direct environment training data, the interaction components (G×E or G×O) may have introduced greater uncertainty into the predictions. Consistent with this, Jarquin et al. (2020) also reported that CV0 is more challenging than CV2 and CV1 because it requires prediction in previously untested environments, thereby limiting the transferability of G×E or covariate relationships.

The final scenario, CV00, was the most challenging scenario because it required predicting new lines in new environments. Prediction accuracies for all four traits were below 0.2, indicating that potential gains from modeling G×E or G×O are minimal when limited information is available for either genotype or environment. This is consistent with previous studies showing that, in CV00, the transferability of genotype-environment and covariate relationship is limited (Ankamah-Yeboah et al., 2020; Jarquín et al., 2017). Across traits, model M2 showed the highest prediction accuracy, which is expected because simpler models may be more stable when there is insufficient information to reliably estimate complex interaction patterns. However, given these low accuracies, differences among models should be interpreted cautiously, and strong conclusions about improvements in prediction accuracy cannot be made.

## 5. CONCLUSION

The southern United States often experiences highly variable growing conditions due to weather differences across wheat-growing regions. Prediction models that incorporate the main and interaction effects of high-dimensional marker data and weather-based responses, such as our M5 model, can significantly improve prediction accuracy. However, the benefit of modelling G×E alone (M3 model) appears to be trait-dependent, especially when the interaction pattern is repeatable even without weather covariates. In contrast, the low prediction accuracies observed when predicting new lines in new environment suggest further improvement will require better multi-environment trial coverage and richer environmental descriptions, including crop growth-stage-specific weather indices.

## Abbreviations

BLUE: best linear unbiased estimate
CV: cross validation
GAWN: gulf Atlantic wheat nursery
GBLUP: genetic best linear unbiased prediction
GBS: genotype-by-sequencing
GEBV: genomic estimated breeding values
GP: genomic prediction
GS: genomic selection
G×E: genotype-by-environment
G×O: genotype-by-environmental covariate
HD: heading date
HT: plant height
IID: independently and identically distributed
MET: multi-environment trials
MSE: mean squared error
NC: North Carolina
PA: Prediction accuracy
PCA: principal component analysis
RNM: reaction norm model
SNP: single nucleotide polymorphisms
SRWW: soft red winter wheat
TW: test weight
Y: grain yield

## AUTHOR CONTRIBUTIONS

**Shailesh Raj Acharya**: Data curation; formal analysis; investigation; methodology; writing– original draft; writing–review and editing. **Julian Garcia Abadillo Velasco:** Formal analysis, writing review and editing. **Jeanette Lyerly:** Data curation, Writing review and editing. **Gina Brown-Guedira:** Writing review and editing. **Diego Jarquin:** Formal analysis, writing review and editing. **Nonoy Bandillo:** Conceptualization; funding acquisition; investigation; methodology; project administration; supervision; writing–review and editing.

## ACKNOWLEDGMENTS

We extend our sincere gratitude to the genotyping team at North Carolina State University (NCSU) and the NC Small Grains Growers Association for their invaluable contributions to this research. This project was supported by Bandillo Hatch Funding (Award No. 04027) at NCSU and the US Wheat and Barley Scab Initiative (Award No. 59-0206-4-046).

## DATA AVAILABILITY STATEMENT

Data are available upon request from the corresponding author.

## REFERENCES

Ankamah-Yeboah, T., Janss, L. L., Jensen, J. D., Hjortshøj, R. L., & Rasmussen, S. K. (2020). Genomic Selection Using Pedigree and Marker-by-Environment Interaction for Barley Seed Quality Traits From Two Commercial Breeding Programs. Frontiers in Plant Science, 11. 10.3389/fpls.2020.00539

Brault, C., Conley, E. J., Read, A. C., Green, A. J., Glover, K. D., Cook, J. P., Gill, H. S., Fiedler, J. D., & Anderson, J. A. (2025). Improving genomic prediction for plant disease using environmental covariates. Plant Methods, 21(1), 114. 10.1186/s13007-025-01418-0

Browning, B. L., & Browning, S. R. (2016). Genotype Imputation with Millions of Reference Samples. American Journal of Human Genetics, 98(1), 116–126. 10.1016/j.ajhg.2015.11.020

Butler, D. G., Cullis, B. R., Gilmour, A. R., Gogel, B. G., & Thompson, R. (2023). ASReml-R Reference Manual Version 4.2. VSN International Ltd.

Grolemund, G., & Wickham, H. (2011). Dates and Times Made Easy with lubridate. Journal of Statistical Software, 40(3), 1–25. 10.18637/jss.v040.i03

Han, L., Wang, X., Benke, R., Tibbs-Cortes, L. E., Zhao, P., Sanguinet, K. A., Zhang, Z., Xu, S., Yu, J., & Li, X. (2025). Integrated phenomic and genomic analyses unveil modes of altered phenotypic plasticity during wheat improvement. Genome Biology, 26(1), 256. 10.1186/s13059-025-03740-1

Jarquín, D., Crossa, J., Lacaze, X., Du Cheyron, P., Daucourt, J., Lorgeou, J., Piraux, F., Guerreiro, L., Pérez, P., Calus, M., Burgueño, J., & de los Campos, G. (2014). A reaction norm model for genomic selection using high-dimensional genomic and environmental data. Theoretical and Applied Genetics, 127(3), 595–607. 10.1007/s00122-013-2243-1

Jarquin, D., Howard, R., Crossa, J., Beyene, Y., Gowda, M., Martini, J. W. R., Covarrubias Pazaran, G., Burgueño, J., Pacheco, A., Grondona, M., Wimmer, V., & Prasanna, B. M. (2020). Genomic Prediction Enhanced Sparse Testing for Multi-environment Trials. G3 Genes|Genomes|Genetics, 10(8), 2725–2739. 10.1534/g3.120.401349

Jarquín, D., Lemes da Silva, C., Gaynor, R. C., Poland, J., Fritz, A., Howard, R., Battenfield, S., & Crossa, J. (2017). Increasing Genomic-Enabled Prediction Accuracy by Modeling Genotype × Environment Interactions in Kansas Wheat. The Plant Genome, 10(2), plantgenome2016.12.0130. 10.3835/plantgenome2016.12.0130

Li, X., Guo, T., Wang, J., Bekele, W. A., Sukumaran, S., Vanous, A. E., McNellie, J. P., Tibbs-Cortes, L. E., Lopes, M. S., Lamkey, K. R., Westgate, M. E., McKay, J. K., Archontoulis, S. V., Reynolds, M. P., Tinker, N. A., Schnable, P. S., & Yu, J. (2021). An integrated framework reinstating the environmental dimension for GWAS and genomic selection in crops. Molecular Plant, 14(6), 874–887. 10.1016/j.molp.2021.03.010

McBreen, J., Babar, Md. A., Jarquin, D., Khan, N., Harrison, S., DeWitt, N., Mergoum, M., Lopez, B., Boyles, R., Lyerly, J., Murphy, J. P., Shakiba, E., Sutton, R., Ibrahim, A., Howell, K., Smith, J. H., Brown-Guedira, G., Tiwari, V., Santantonio, N., & Van Sanford, D. A. (2025). Enhancing prediction accuracy of grain yield in wheat lines adapted to the southeastern United States through multivariate and multi-environment genomic prediction models incorporating spectral and thermal information. The Plant Genome, 18(1), e20532. 10.1002/tpg2.20532

Puglisi, D., Crossa, J., Cuevas, J., Fania, F., Vitale, P., & De Vita, P. (2026). Multi-trait and multi-environment genomic prediction enhances yield components improvement in durum wheat. Frontiers in Plant Science, 17. 10.3389/fpls.2026.1759897

Sparks, A. H. (2018). nasapower: A NASA POWER Global Meteorology, Surface Solar Energy and Climatology Data Client for R. Journal of Open Source Software, 3(30), 1035. 10.21105/joss.01035

Sukumaran, S., Jarquin, D., Crossa, J., & Reynolds, M. (2018). Genomic-enabled Prediction Accuracies Increased by Modeling Genotype × Environment Interaction in Durum Wheat. The Plant Genome, 11(2), 170112. 10.3835/plantgenome2017.12.0112

USDA. (2025). USDA/NASS 2024 State Agriculture Overview for North Carolina. https://www.nass.usda.gov/Quick_Stats/Ag_Overview/stateOverview.php?state=NORTH+CAROLINA

Winn, Z. J., Lyerly, J. H., Brown-Guedira, G., Murphy, J. P., & Mason, R. E. (2023). Utilization of a publicly available diversity panel in genomic prediction of Fusarium head blight resistance traits in wheat. The Plant Genome, 16(3), e20353. 10.1002/tpg2.20353

