## Supplemental for "Reaction Norm Modeling of High-Dimensional Genomic and Environmental Data Improves Prediction Accuracy in Winter Wheat"

Pages: 9
Supplemental Tables: 4
Supplemental Figures: 4

#### Supplemental tables

**Supplemental table S1.** Broad sense heritability ($H^2$) estimation across all environments for yield, test-weight, heading date and plant height. Heritability estimates are only reported where the available data supported reliable estimation. Entries left blank denote cases in which $H^2$ could not be obtained due to insufficient data, limited replication or unsuccessful model fitting.

| Environment | Yield | TestWt | HeadDate | PlantHt |
|---|---|---|---|---|
| CLAYTON2017 | 0.70 | 0.87 | 0.95 | 0.33 |
| CLAYTON2018 | 0.62 | 0.77 | | 0.59 |
| CLAYTON2019 | 0.33 | 0.81 | | |
| CLAYTON2020 | | 0.08 | | |
| CLAYTON2021 | 0.83 | 0.95 | | |
| CLAYTON2022 | 0.61 | 0.96 | | |
| KINSTON2017 | 0.60 | 0.21 | 0.86 | |
| KINSTON2018 | 0.66 | 0.32 | 0.83 | |
| KINSTON2019 | 0.47 | 0.48 | 0.92 | 0.81 |
| KINSTON2020 | 0.70 | 0.83 | 0.88 | |
| KINSTON2021 | | 0.33 | 0.87 | |
| KINSTON2022 | 0.61 | 0.82 | 0.89 | |
| KINSTON2023 | 0.70 | 0.91 | 0.35 | |
| PLYMOUTH2020 | 0.64 | 0.84 | | |
| PLYMOUTH2021 | 0.60 | 0.43 | | |
| PLYMOUTH2022 | 0.86 | 0.67 | | |
| RALEIGH2017 | 0.87 | 0.90 | | 0.75 |
| RALEIGH2019 | 0.53 | 0.91 | 0.87 | 0.78 |
| RALEIGH2020 | 0.73 | 0.92 | 0.96 | |
| RALEIGH2021 | | | | |
| SALISBURY2017 | 0.82 | 0.94 | | |
| SALISBURY2019 | 0.80 | 0.78 | | 0.89 |
| SALISBURY2020 | 0.63 | 0.89 | | 0.88 |
| SALISBURY2021 | 0.56 | 0.85 | | |

**Supplemental table S2.** Latitude and longitude of coordinates of four field locations involved in this study.

| Location | Latitude (ºN) | Longitude (ºW) |
|---|---|---|
| Clayton, NC | 35.67 | 78.49 |
| Kinston, NC | 35.29 | 77.57 |
| Salisbury, NC | 35.69 | 80.62 |
| Plymouth, NC | 35.87 | 76.66 |

**Supplemental table S3.** Variance partitioning across all five models for four traits. The abbreviation varE, varL, varG, varGE, varGO and varR represent variance for environment, line, genomic, genotype-by-environment interaction, genotype-by-weather covariate interaction and residuals respectively.

| | | varE | varL | varG | varGE | varGO | varR |
|---|---|---|---|---|---|---|---|
| **Head Date** | M1(E+L) | 78.7 | 6.8 | | | | 14.5 |
| | M2(E+L+G) | 76.8 | 1.9 | 9.6 | | | 11.8 |
| | M3(E+L+G+GE) | 77.1 | 1.7 | 6.3 | 9.1 | | 5.7 |
| | M4(E+L+G+GO) | 77.3 | 1.5 | 7.3 | | 6.9 | 7.0 |
| | M5(E+L+G+GE+GO) | 78.0 | 1.4 | 6.2 | 6.1 | 2.9 | 5.4 |

| | | | | | | | |
|---|---|---|---|---|---|---|---|
| **PlantHt** | M1(E+L) | 46.1 | 39.2 | | | | 14.8 |
| | M2(E+L+G) | 47.6 | 18.2 | 20.5 | | | 13.6 |
| | M3(E+L+G+GE) | 40.7 | 15.2 | 26.9 | 8.9 | | 8.3 |
| | M4(E+L+G+GO) | 43.7 | 12.4 | 25.1 | | 6.3 | 12.4 |
| | M5(E+L+G+GE+GO) | 47.4 | 12.7 | 21.7 | 7.2 | 4.6 | 6.3 |

| | | | | | | | |
|---|---|---|---|---|---|---|---|
| **TestWt** | M1(E+L) | 55.4 | 22.2 | | | | 22.3 |
| | M2(E+L+G) | 51.1 | 7.8 | 19.6 | | | 21.5 |
| | M3(E+L+G+GE) | 51.2 | 9.1 | 16.5 | 10.6 | | 12.5 |
| | M4(E+L+G+GO) | 48.7 | 6.7 | 17.3 | | 10.9 | 16.4 |
| | M5(E+L+G+GE+GO) | 49.2 | 8.7 | 16.3 | 7.0 | 5.9 | 12.9 |

| | | | | | | | |
|---|---|---|---|---|---|---|---|
| **Yield** | M1(E+L) | 55.9 | 14.3 | | | | 29.8 |
| | M2(E+L+G) | 54.8 | 3.4 | 13.2 | | | 28.6 |
| | M3(E+L+G+GE) | 54.1 | 3.6 | 11.2 | 14.6 | | 16.5 |
| | M4(E+L+G+GO) | 52.0 | 2.7 | 10.1 | | 13.7 | 21.4 |
| | M5(E+L+G+GE+GO) | 53.5 | 3.0 | 10.5 | 10.6 | 5.9 | 16.4 |

**Supplemental table S4.** Prediction accuracies across all five models under each cross-validation schemes estimated for all four traits. The abbreviations E, L, G, GE and GO represent environment, line, genomic, genotype-by-environment interaction and genotype-by-weather covariate interaction respectively.

| | MODELS | CV00 | CV0 | CV1 | CV2 |
|---|---|---|---|---|---|
| YIELD | M1 (E + L) | 0.003 | 0.268 | -0.114 | 0.360 |
| | M2 (E + L+ G) | 0.204 | 0.310 | 0.333 | 0.440 |
| | M3 (E + L + G + GE) | 0.182 | 0.277 | 0.411 | 0.500 |
| | M4(E + L + G + GO) | 0.184 | 0.290 | 0.390 | 0.491 |
| | M5(E +L+G+GE+GO) | 0.172 | 0.282 | 0.413 | 0.504 |
| PLANTHT | M1 (E + L) | -0.032 | 0.430 | -0.263 | 0.433 |
| | M2 (E + L+ G) | 0.102 | 0.463 | 0.137 | 0.504 |
| | M3 (E + L + G + GE) | 0.107 | 0.458 | 0.055 | 0.504 |
| | M4(E + L + G + GO) | 0.037 | 0.422 | 0.143 | 0.490 |
| | M5(E +L+G+GE+GO) | 0.140 | 0.402 | 0.134 | 0.504 |
| TESTWT | M1 (E + L) | 0.001 | 0.311 | -0.093 | 0.411 |
| | M2 (E + L+ G) | 0.156 | 0.340 | 0.275 | 0.461 |
| | M3 (E + L + G + GE) | 0.139 | 0.314 | 0.376 | 0.540 |
| | M4(E + L + G + GO) | 0.157 | 0.347 | 0.355 | 0.530 |
| | M5(E +L+G+GE+GO) | 0.150 | 0.337 | 0.378 | 0.548 |
| HEADDATE | M1 (E + L) | 0.000 | 0.091 | -0.101 | 0.309 |
| | M2 (E + L+ G) | 0.141 | 0.185 | 0.449 | 0.512 |
| | M3 (E + L + G + GE) | 0.095 | 0.186 | 0.493 | 0.555 |
| | M4(E + L + G + GO) | 0.116 | 0.135 | 0.487 | 0.551 |
| | M5(E +L+G+GE+GO) | 0.090 | 0.141 | 0.488 | 0.553 |

### Supplemental figures

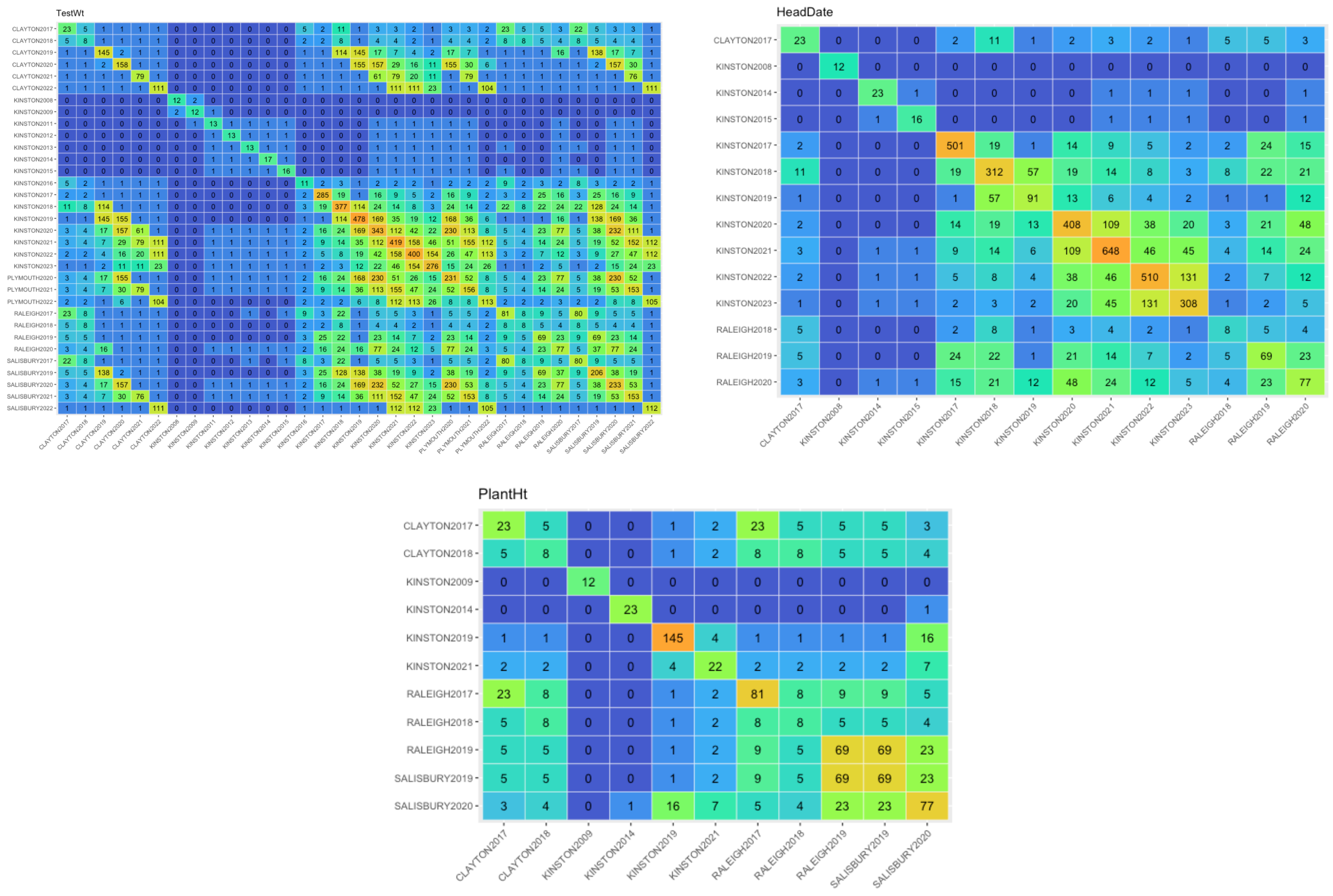

**Supplemental figure S1.** Total number of overlapping genotypes between environments for *'TestWt', HeadDate and Plant Height.* The 33x33 matrix shows the number of lines in each environment (location-year combinations) on the diagonal. The upper off-diagonal shows common lines between environments, while the lower off-diagonal shows lines not shared between environments.

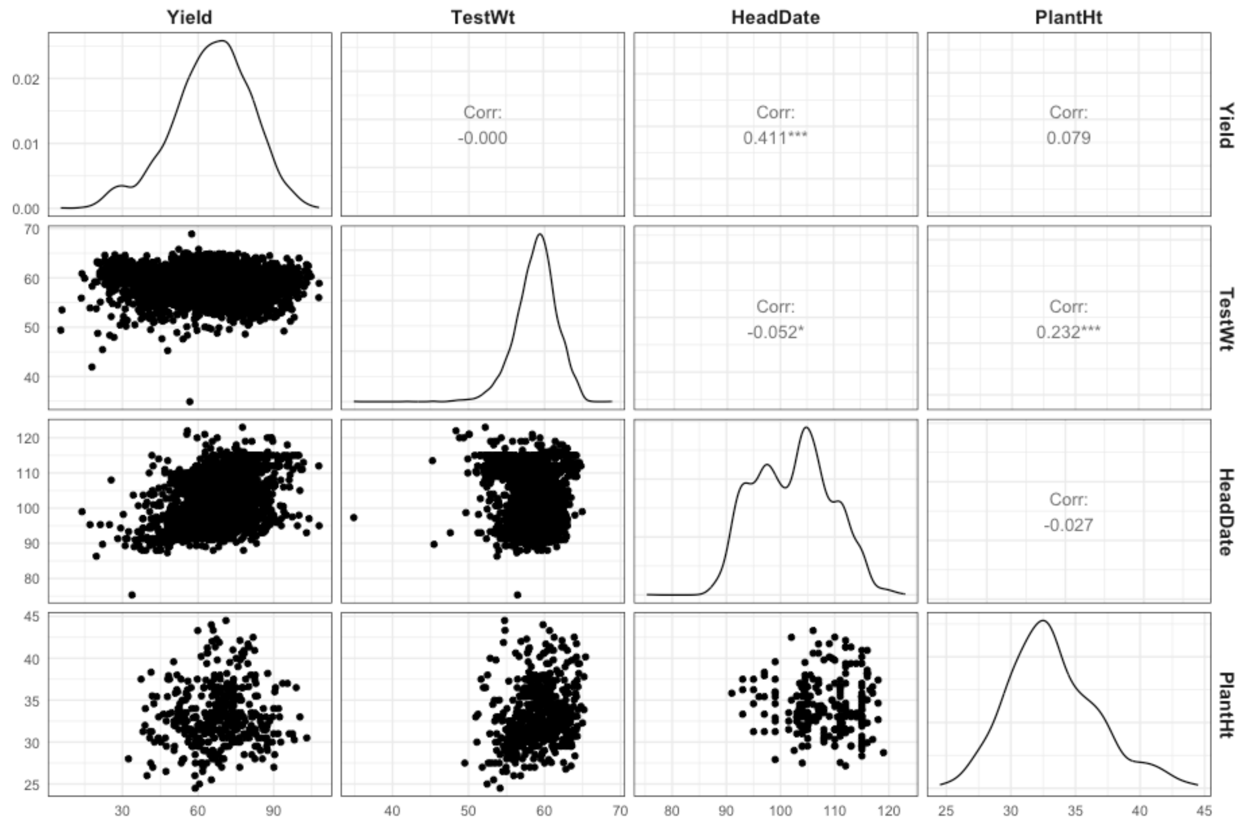

**Supplemental figure S2.** Genetic correlation between traits. Scatterplot matrix showing relationship among yield, test-weight, heading date and plant height. The diagonal panels represent the distribution of each variable as density curves. The lower triangular panel shows pairwise scatterplots and the upper panel represent the corresponding correlation coefficients between trait pairs.

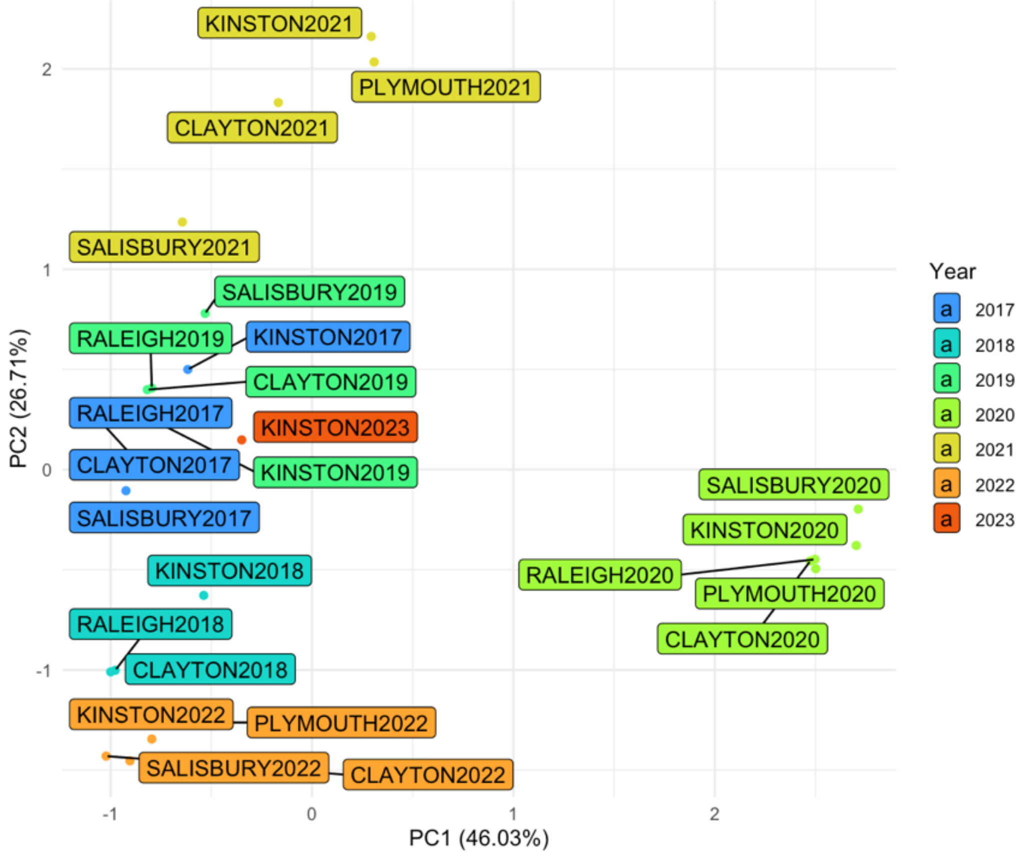

**Supplemental figure S3.** PCA derived from NASA weather data. Color denotes the year associated with the environment. The first 2 PC explained 46.03% and 26.71% of the variance, respectively. 80% and 95% of the variance was explained with 3 and 5 PCs, respectively.

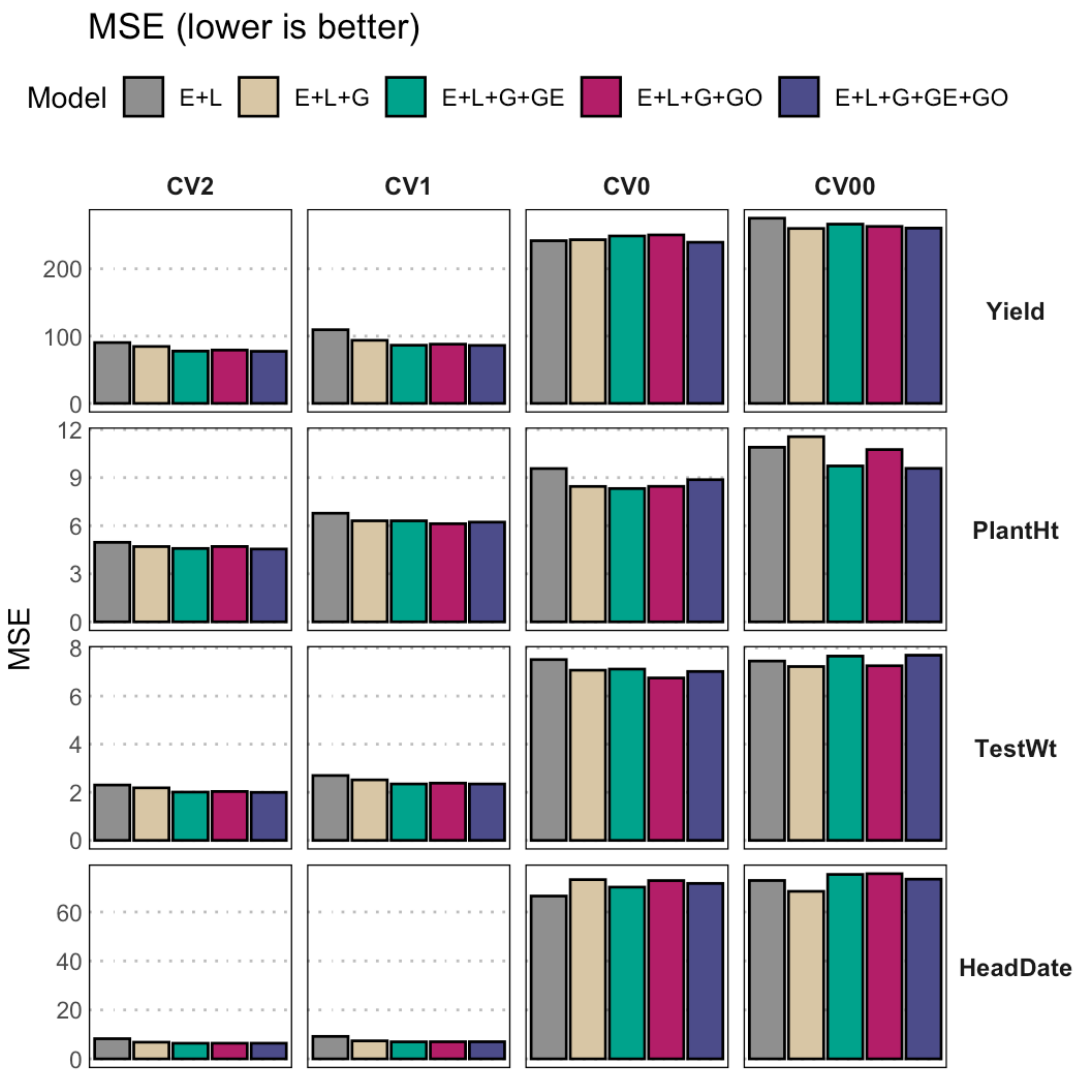

**Supplemental figure S4.** Mean Squared Error (MSE) achieved by the models under different cross-validation scenarios (columns) and predicting different traits (rows). In each grid, MSE is denoted in the y-axis, while models are denoted in the x-axis and color. The y-axis is scaled for each trait.
